# MYB28 and MYB29 transcription factors regulate iron homeostasis and iron-mobilizing coumarin biosynthesis in *Arabidopsis thaliana*

**DOI:** 10.64898/2026.09.09.750326

**Authors:** Agustín J. Marín-Peña, Inmaculada Coleto, José Alberto Urbano-Gámez, Alice Rossille, Diego Tazueco, Sunshuke Watanabe, Daniela Salazar-Gutierrez, Antonio Santiago, Meng-Bo Tian, Hannetz Roschzttardtz, Joaquín Medina, Jose Tomás Matús, Christian Dubos, Daniel Marino

## Abstract

Iron (Fe) deficiency is a major constraint for plant growth and triggers extensive physiological and transcriptional reprogramming to maintain Fe homeostasis. Here, we identify the glucosinolate-associated transcription factors MYB28 and MYB29 as previously unrecognized regulators of *Arabidopsis thaliana* adaptation to Fe deficiency. Across various growth systems, loss of MYB28 increased sensitivity to Fe deficiency, whereas the *myb28myb29* double mutant displayed stronger chlorosis, reduced root growth and impaired biomass accumulation, indicating cooperative but unequal functions of these transcription factors. Despite their enhanced Fe-deficiency phenotype, double mutant plants accumulated higher Fe levels in roots and exhibited stronger induction of canonical Fe-deficiency responses, suggesting impaired Fe utilization or distribution. RNA-seq revealed extensive transcriptional reprogramming under Fe deficiency, with pronounced deregulation of genes involved in Fe homeostasis, redox processes and growth, particularly in the double mutant. Among these, *SCOPOLETIN 8-HYDROXYLASE* (*S8H*) emerged as a major target gene of MYB28. The expression of *S8H* was almost abolished in *myb28* and *myb28myb29* mutants, whereas expression of other coumarin biosynthetic genes remained largely unaffected. Promoter activation assays demonstrated that MYB28 activates the *S8H* promoter, and metabolic analyses showed accumulation of scopolin together with reduced fraxin levels in the mutants, consistent with impaired S8H activity. Collectively, our results identify MYB28 as a key regulator linking specialized metabolism to Fe homeostasis through control of coumarin biosynthesis, thereby expanding the biological functions of MYB28 and MYB29 transcription factors.

## Introduction

Iron (Fe) is an essential micronutrient for all living organisms. Although Fe is the fourth most abundant element on Earth, plants frequently experience Fe deficiency in nearly one-third of cultivated land. This issue is due to the relationship between soil pH and Fe availability. At neutral or alkaline pH, Fe is predominantly present as insoluble oxides and hydroxides, which significantly limit its uptake by plants. However, as soil pH decreases, Fe becomes more accessible in the form of soluble ferrous (Fe^2+^) or ferric (Fe^3+^) ions (Guerinot & Yi, 1994). Due to its dual role as both an electron donor and acceptor, Fe functions as a cofactor for many enzymes, playing a critical role in key physiological processes such as respiration, sulfur (S) and nitrogen (N) assimilation (Touraine et al., 2019; Przybyla-Toscano et al., 2021). Consequently, Fe deficiency can have serious consequences on crop performance, including chlorosis, stunted vegetative growth, and substantial losses in crop yield and quality.

To efficiently acquire Fe, dicotyledonous and non-grass monocot species secrete protons into the rhizosphere to increase the solubility of Fe^3+^, which is then reduced to Fe²⁺ by ferric reductases and taken up into root cells by specific Fe transporters. In Arabidopsis, this three-step process is mediated by the proton ATPase AHA2 (AUTOINHIBITED PLASMA MEMBRANE H^+^-ATPase 2), FRO2 (FERRIC REDUCTION-OXIDASE 2) and the IRT1 (IRON-REGULATED TRANSPORTER 1), respectively. In contrast, grasses, secrete phytosiderophores that chelate Fe³⁺ and facilitate its absorption in the form of Fe³⁺-phytosiderophore complexes (Kobayashi et al., 2019).

Besides the classical Fe-reduction strategy, dicots are also capable of secreting phenolic and/or flavin-type metabolites that facilitate Fe acquisition (Rodríguez-Celma et al., 2013). In particular, under limited Fe availability, Arabidopsis roots synthesize and secrete Fe-mobilizing coumarins (FMCs) into the rhizosphere. FMCs are phenylpropanoid-derived compounds that contribute to solubilize and reduce Fe^3+^ (Robe et al., 2021a). In addition, they may form complexes with Fe^3+^ that can be taken up by root cells (Robe et al., 2025). In Arabidopsis, the biosynthesis of FMCs begins with F6′H1 (FERULOYL-COA 6′-HYDROXYLASE 1), which catalyses the ortho-hydroxylation of feruloyl-CoA to produce 6-hydroxyferuloyl-CoA (Schmidt et al., 2014). This intermediate is subsequently converted into scopoletin by the acyltransferase COSY (COUMARIN SYNTHASE) (Vanholme et al., 2019). Because scopoletin lacks a catechol moiety, it is unable to bind Fe and, therefore, is not directly involved in Fe acquisition (Rajniak et al., 2018). The two primary FMCs, fraxetin and sideretin, are biosynthesized by S8H (SCOPOLETIN 8-HYDROXYLASE), a 2-oxoglutarate-dependent dioxygenase (Siwinska et al., 2018; Tsai et al., 2018), and CYP82C4, a P450-dependent monooxygenase, respectively. Finally, FMCs, as well as scopolotein, are secreted into the rhizosphere through the ABCG transporter PDR9 (Pleiotropic Drug Resistance 9) (Fourcroy et al., 2014) or stored into the vacuoles following their glycosylation by specific UDP-GLYCOSYLTRANSFERASE such as UGT72E1, E2 and E3 (Wu et al., 2022), in the form of fraxin, sideretin glycoside or scopolin, respectively. Scopoletin secretion is further regulated by the deglycosylation of scopolin, a process facilitated by BGLU42 (β GLUCOSIDASE 42) (Zamioudis et al., 2014).

Plant response to Fe deficiency is tightly regulated by a complex transcriptional regulatory cascade that ensures Fe availability while preventing Fe toxicity. In this cascade, bHLH transcription factors (TFs) play a predominant role, with more than 15 members identified in Arabidopsis (Gao & Dubos, 2024). Briefly, *AHA2*, *FRO2* and *IRT1* expression is induced in response to Fe deficiency by bHLH complexes comprising FIT/bHLH29 (FER-LIKE IRON DEFICIENCY INDUCED TRANSCRIPTION FACTOR) and bHLHs from clade Ib (i.e. bHLH38, bHLH39, bHLH100 and bHLH101). The expression of these bHLH TFs is also regulated by another set of bHLHs belonging to clade IVb (i.e., bHLH11, PYE/bHLH47: POPEYE, and URI/bHLH121: UPSTREAM REGULATOR OF IRT1) and clade IVc (IDT1/bHLH34: IRON DEFICIENCY TOLERANT1, bHLH104, ILR3/bHLH105: IAA-LEUCINE RESISTANT 3 and bHLH115). The stability of FIT, URI, ILR3 and bHLH115 is regulated by specific E3-ubiquitin ligases (i.e., BTS/BRUTUS, BTSL1/BRUTUS LIKE 1 and BTSL2/BRUTUS LIKE 2) whose interaction leads to their degradation via the 26S proteasome (Selote et al., 2025; Rodríguez-Celma et al., 2019; Zhao et al., 2026). The activity of these E3-ubiquitin ligases is itself inhibited via their interaction with peptides from the IMA/FEP (IRONMAN/ FE-UPTAKE-INDUCING PEPTIDE) family (Grillet et al., 2018; Hirayama et al., 2018). In addition, the Fe-deficiency regulatory network also comprises TFs from other families such as ARF, bZIP, EIL, C2H2, NF-Y, MYB or WRKY. Indeed, this signalling network regulates not only Fe uptake, but also Fe storage and distribution (Gao & Dubos 2021; Riaz & Guerinot 2021). For instance, MYB72 was shown to be involved in the regulation of the biosynthesis of both nicotianamine and coumarins that are required for Fe phloem transport and Fe uptake, respectively (Palmer et al., 2013; Zamioudis et al., 2014).

MYB28 and MYB29 are positive regulators of the aliphatic glucosinolate (GSL) biosynthetic pathway in Brassicaceae, playing major roles in the regulation of long- and short-chain aliphatic GSLs, respectively (Beekwilder et al., 2008; Sønderby et al., 2007). Consistent with this function, the *myb28myb29* double mutant exhibits an almost complete absence of aliphatic GSLs. The GSL-dependent function of MYB28 and MYB29 has also been associated with additional phenotypes, including drought responses, as the *myb28myb29* exhibits drought sensitivity (Salehin et al., 2019).

Although both TFs cooperate in the regulation of GSL biosynthesis, they also display non-additive effects on the transcriptional regulation of other pathways (Sønderby et al., 2010). For example, MYB28 contributes to S homeostasis by directly regulating S seed storage proteins through binding to the *At2S4* (*SEED STORAGE ALBUMIN 4*) promoter (Aarabi et al., 2016). Regarding MYB29, Gaudinier et al., (2018) reported a nitrate-dependent root-architecture phenotype in *myb29-1* mutant, while Zhang et al., (2017) identified MYB29 as an indirect negative regulator of *AOX1a (ALTERNATIVE OXIDASE 1a*) expression. However, these two studies did not evaluate whether these functions were associated with GSL metabolism.

Furthermore, Coleto et al. (2021) reported a GLS-independent function of MYB28 and MYB29 under ammonium (NH_4_ ^+^) stress. One of the causes of NH_4_ ^+^ toxicity is associated with altered Fe homeostasis (Liu et al., 2022; Coleto et al., 2023). Gene expression analyses revealed that, under NH_4_ ^+^ stress, the *myb28myb29* mutant exhibited altered expression of several Fe-related genes. Moreover, the sensitivity of *myb28myb29* to NH_4_ ^+^ was prevented by the supply of a higher Fe concentration to the media (Coleto et al., 2021).

In this context, we aimed at exploring the potential function of MYB28 and MYB29 as regulators of Fe nutrition in *Arabidopsis thaliana*. Notably, we report a novel function for these two TFs in the regulation of Fe-deficiency responses, through the direct regulation of the FMC biosynthesis pathway.

## Results

### myb28 and myb28myb29 are hypersensitive to Fe deficiency

In Coleto et al. (2021) we reported that the absence of both MYB28 and MYB29 TFs leads to NH_4_^+^ hypersensitivity, a phenotype that was associated with NH_4_^+^-mediated alterations of Fe homeostasis. In the present work, to explore whether MYB28 and MYB29 TFs play a direct role in the regulation of Fe homeostasis, we evaluated their involvement in Arabidopsis response to Fe deficiency.

As Fe is essential for chlorophyll synthesis, leaf chlorosis is one of the most common symptoms of Fe deficiency. After 10 days of Fe deprivation under hydroponic conditions, both *myb28* and *myb28myb29* mutants exhibited enhanced chlorosis compared with WT plants. In contrast, the *myb29* mutant did not show significant differences respect to WT plants (Figure 1A-B). Regarding whole plant biomass accumulation, no significant differences were observed among genotypes or between treatments (Figure 1C).

**Figure 1.**
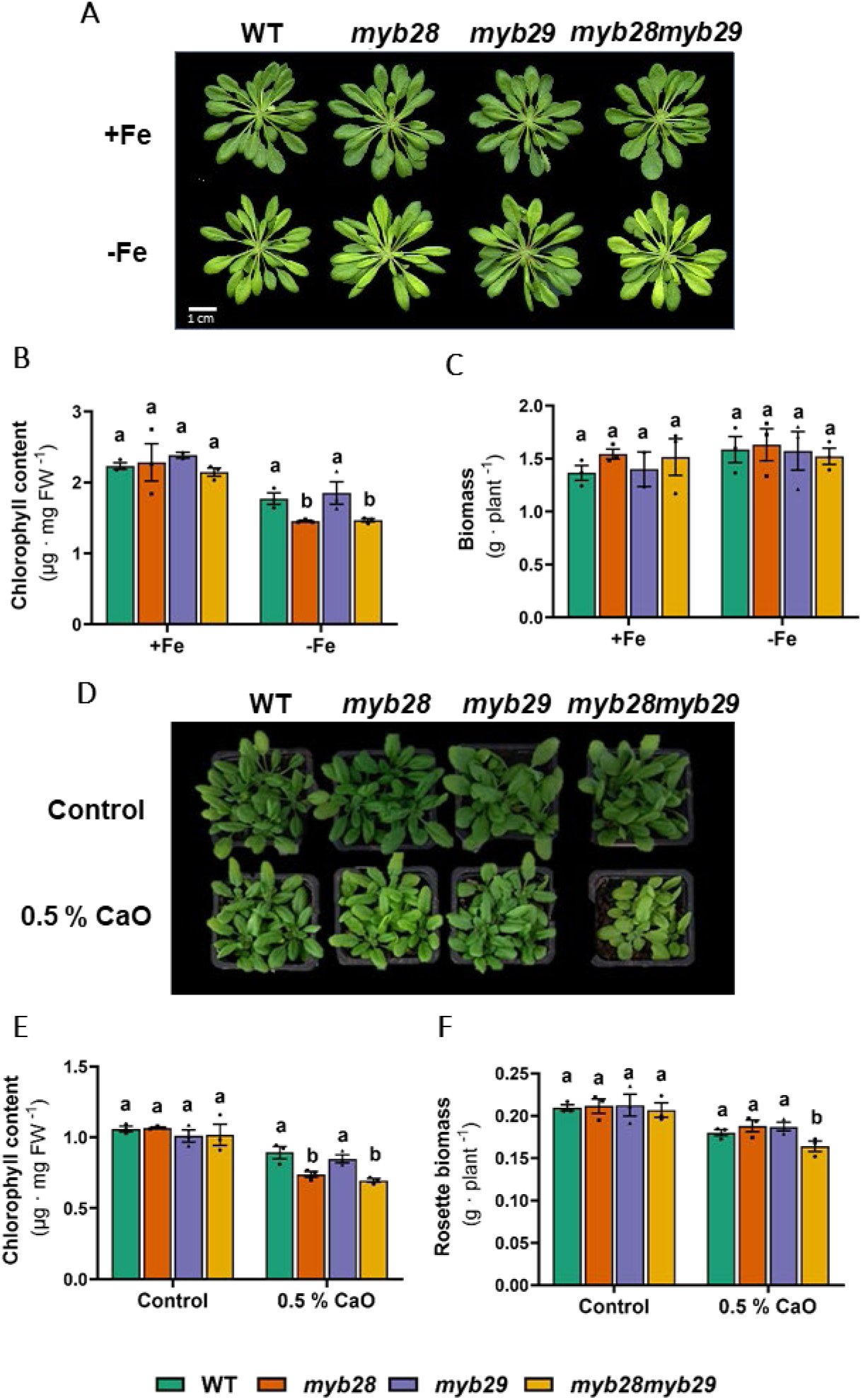
*myb28* and *myb28myb29* plants are hypersensitive to Fe deficiency. (A) Representative images, (B) chlorophyll content and (C) plant biomass of wild-type (WT), *myb28*, *myb29* and *myb28myb29* plants grown hydroponically and subjected to a 10-day exposure to control (+Fe) or Fe-deficiency (-Fe) conditions. (B-C) Bars represent mean ± SEM of a representative experiment (n = 3, each replicate consists of a pool of 5 plants grown in the same hydroponic tank). (D) Representative images, (E) chlorophyll content and (F) rosette biomass of WT, *myb28*, *myb29* and *myb28myb29* plants grown in peat under control conditions or in presence of 0.5 % CaO. (E-F) Values represent mean ± SEM of a representative experiment (n = 3, each replicate consists of a pool of 5 plants grown in the same pot). At least three independent experiments showing similar results were performed. Different letters indicate significant differences among genotypes (One-way ANOVA followed by Duncan’s *post hoc* test, P ≤ 0.05).

To further investigate the effects of long-term Fe limitation, plants were grown in peat supplemented with 0.5% CaO, thereby exposing them to reduced Fe availability from germination onwards. Under this condition, chlorophyll content followed the same pattern as observed in the hydroponics, with *myb28* and *myb28myb29* showing stronger chlorosis than WT plants, whereas *myb29* did not differ from the WT (Figure 1 D-E). Fe deficiency resulted in lower rosette biomass accumulation. Importantly, the *myb28myb29* double mutant displayed a greater reduction in biomass compared with WT plants or the individual *myb28* and *myb29* mutants (Figure 1F). Altogether, these results provide clear evidence for the involvement of MYB28 and MYB29 in the Arabidopsis response to Fe deficiency, with MYB28 playing a predominant role.

In order to characterize the phenotype of these lines under Fe deficiency and to assess the impact of this treatment on root elongation, we performed *in vitro* experiments, exposing plants to decreasing concentrations of Fe, ranging from optimal level (100 µM) to complete deficiency (0 µM) from germination. Phenotypic analyses confirmed the results obtained in the hydroponic- and CaO-based experiments. The *myb28myb29* double mutant exhibited reduced biomass and root elongation when Fe availability was below 10 µM, while chlorosis became evident below 5 µM (Figure 2A-C and S1). The *myb28* mutant did not show significant differences in biomass regardless of Fe concentration, although its chlorotic phenotype was comparable to that of the double mutant. The *myb29* mutant did not exhibit symptoms of hypersensitivity to Fe deficiency but contributed to the enhanced sensitivity to Fe-deficiency observed in the *myb28myb29* double mutant (Figure 2A-C and S1).

**Figure 2.**
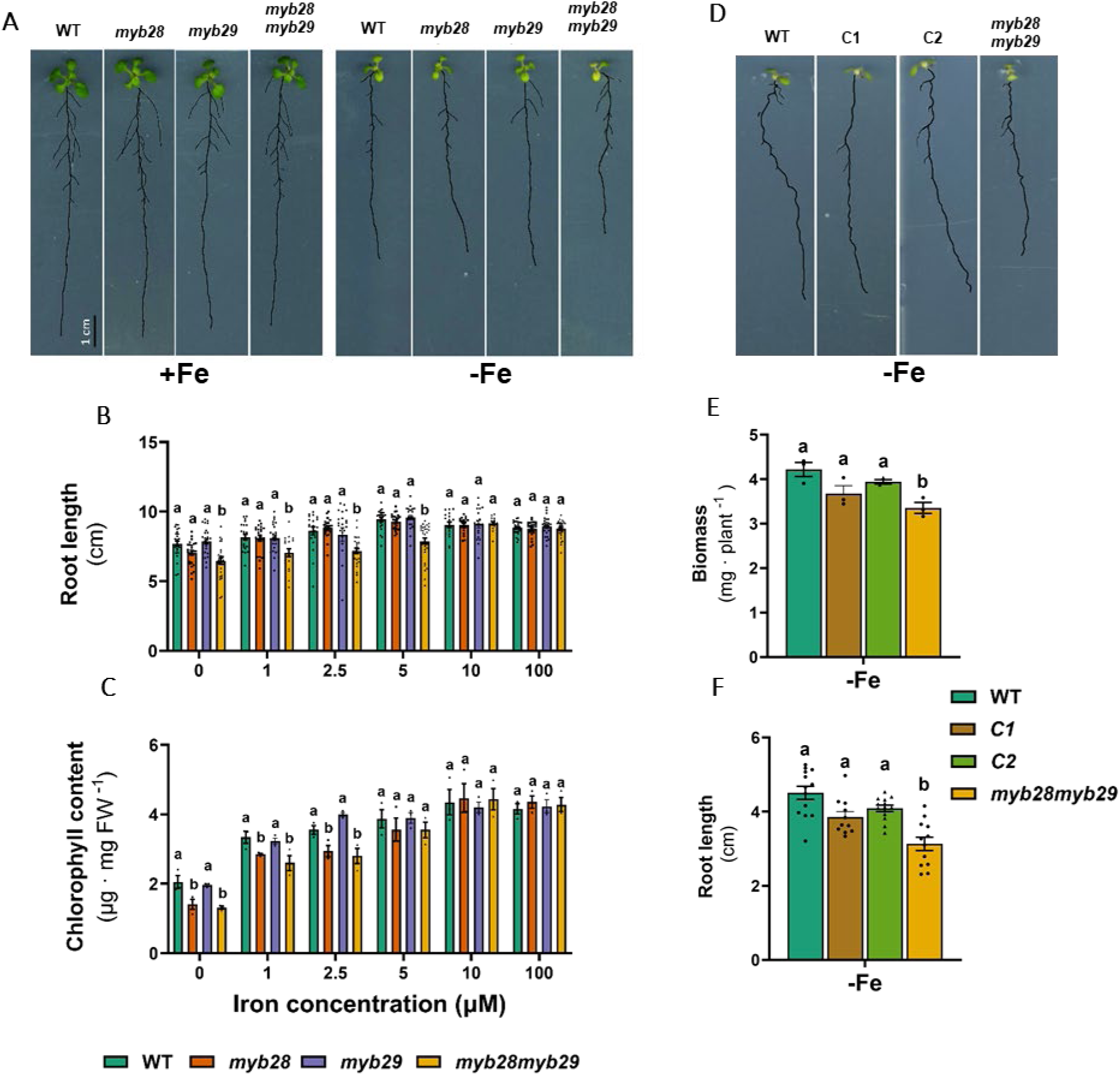
*myb28* and *myb28myb29* display a different degree of sensitivity to Fe deficiency. (A) Representative images, (B) root length and (C) chlorophyll content of WT, *myb28*, *myb29* and *myb28myb29* plants grown *in vitro* for 15 days under control (+Fe) or Fe deficiency (-Fe) conditions. (B-C) Bars represent mean ± SEM (For root length, n = 20–26; each replicate consisted of an individual plant. For chlorophyll, n = 3; each replicate consists of a pooled sample of 20 plants. (D) Representative image, (E) plant biomass and (F) primary root length of WT, *myb28myb29* and two independent lines (C1 and C2) of *myb28myb29* complemented with *MYB28* (*pMYB28:gMYB28:GFP*). Plants were cultured *in vitro* for 10 days under Fe deficiency conditions (-Fe). (E-F) Bars represent mean ± SEM (for root length n = 25, each replicate consists of an individual plant; for biomass n = 3, each replicate consists of a pool of 10 plants). Different letters indicate significant differences within each treatment (One-way ANOVA followed by Duncan’s *post hoc* test, P ≤ 0.05). Three independent experiments showing similar results were performed.

To confirm the observed phenotypes, we introduced a GFP-tagged genomic *MYB28* construct driven by its native promoter (i.e., *pMYB28:gMYB28*) into the *myb28myb29* double mutant. Two independent transgenic lines exhibiting *MYB28* expression levels similar to WT plants were selected (Figure S2). Both lines restored the growth defects of *myb28myb29* plants under Fe deficiency (Figure 2D-F), confirming that loss of MYB28 activity underlies the observed growth phenotype.

### Fe-deficiency responses are enhanced in myb28 and myb28myb29 mutants

As markers of Fe deficiency, we next analyzed the mRNA abundance of a set of key genes encoding regulatory proteins and peptides involved in Fe-deficiency signaling pathway (Fig 3 and S3). These measurements were performed in seedlings grown *in vitro* exposed to Fe deficiency during 15 days from germination. RT-qPCR experiments revealed that the expression of *ILR3* and *bHLH104* was induced in all three mutants tested when compared to the WT. An induction of the expression was also observed for *bHLH11*, *bHLH38*, *bHLH101* and *MYB72* in both *myb28* and *myb28myb29* mutants, and for *FIT* solely in *myb28myb29*. By contrast, the expression levels of *bHLH34*, *bHLH39*, *bHLH100* and *PYE* in the mutants were similar to those of WT. The induction of *BTS*, *BTSL1*, *BTSL2*, *IMA1*, *IMA2* and *IMA3* expression in response to Fe deficiency was also induced in the mutants compared with the WT. Taken together, these observations indicate that MYB28, together with MYB29, acts as a repressor of the Fe deficiency signaling pathway.

**Figure 3.**
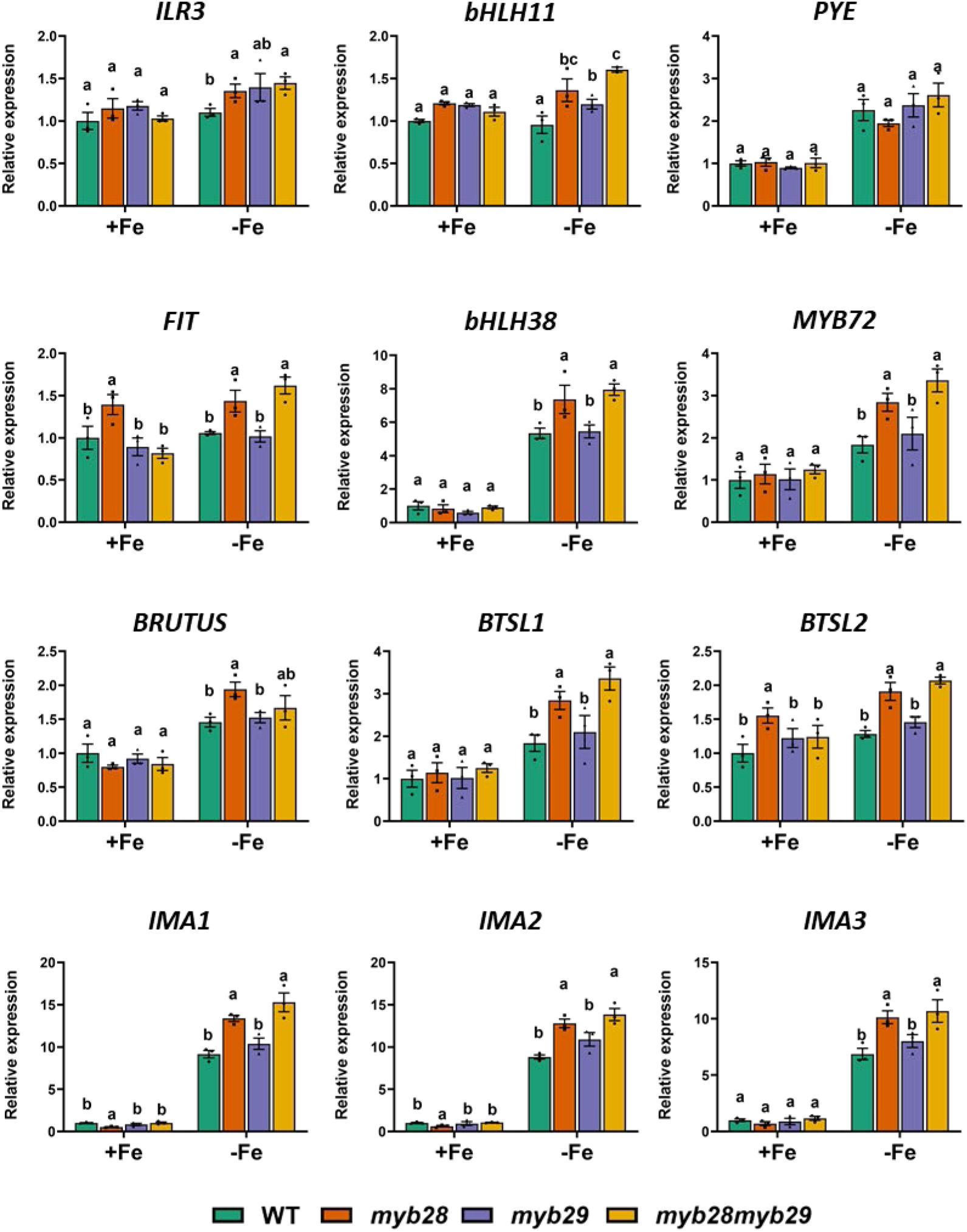
The transcriptional response to Fe deficiency is altered in *myb28* and *myb28myb29*. Expression level of Fe homeostasis-related genes in roots of WT, *myb28*, *myb29* and *myb28myb29* plants grown *in vitro* for 15 days under control (+Fe) or Fe deficiency (-Fe) conditions. Bars represent mean ± SEM (n = 3, each replicate consists of a pool of 12 roots). Different letters indicate significant differences within each treatment (One-way ANOVA followed by Duncan’s post hoc test, P ≤ 0.05). Three independent experiments showing similar results were performed.

We then investigated how loss-of-function of MYB28 and MYB29 influences the expression of genes involved in Fe uptake. As expected, we first observed that the induction of the expression of *FRO2* and *IRT1* (Figure 4A) was similar to that of *FIT* and *bHLH38* (Figure 3). We then investigate the expression of genes involved in coumarin biosynthesis and transport. This analysis revealed that the expression of *F6’H1*, *PDR9* and *BGLU42* was induced in both *myb28* and *myb28myb29* mutant when compared to *myb29* and WT plants (Figure 4B). Strikingly, the expression of *S8H* was nearly abolished in *myb28* and *myb28myb29*, suggesting that both MYBs might regulate fraxetin biosynthesis. Collectively, these results indicate that Fe-deficiency signaling remains functional in all genotypes, while suggesting that *myb28* and *myb28myb29* plants experience an enhanced perception of Fe deficiency.

**Figure 4.**
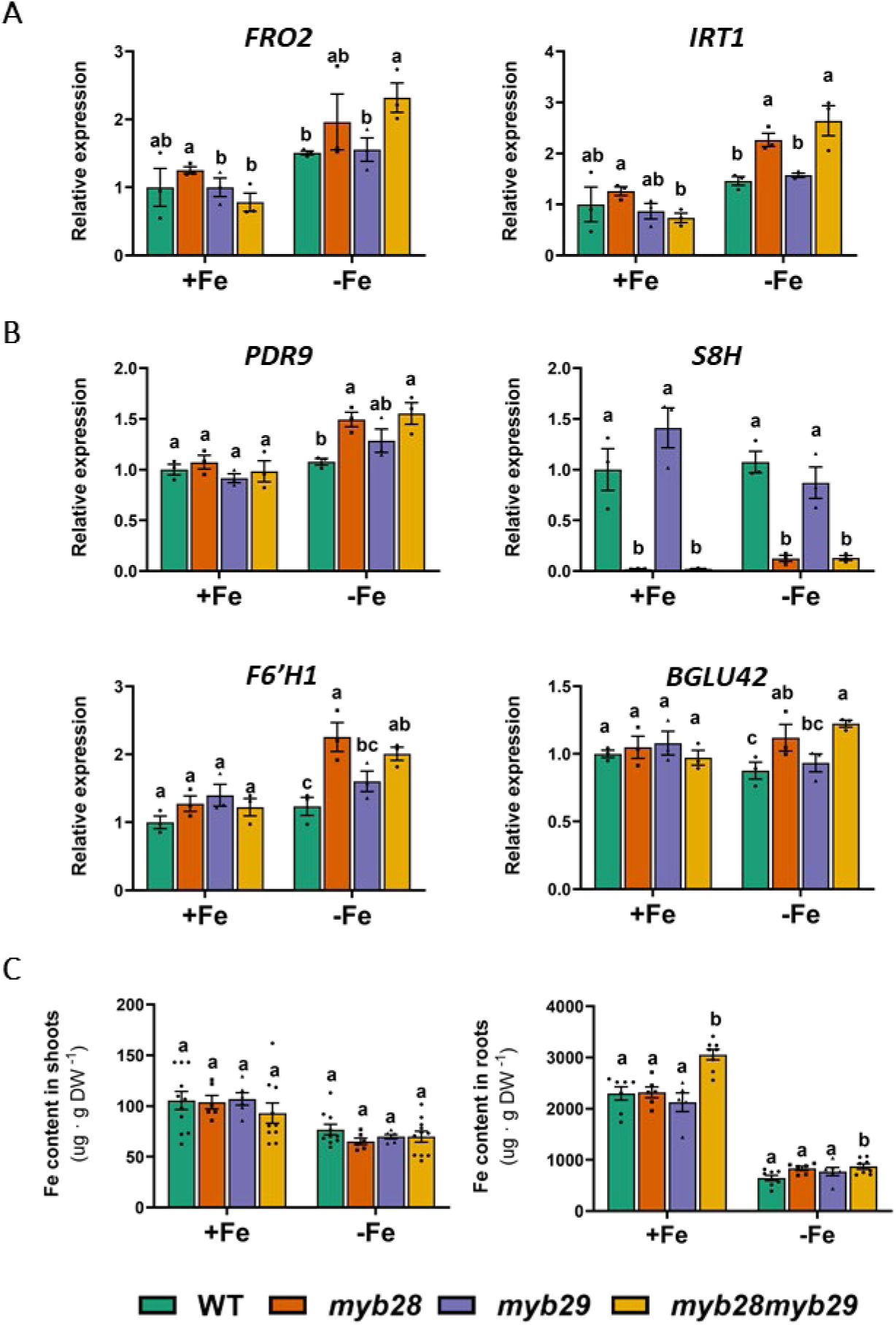
Fe uptake is promoted in *myb28myb29.* (A) Expression level of *FRO2* and *IRT1* genes and (B) of the coumarin-related genes *PDR9*, *S8H*, *F6’H1* and *BGLU42* in roots of WT, *myb28*, *myb29* and *myb28myb29* plants grown *in vitro* for 15 days under control (+Fe) or Fe deficiency (-Fe) conditions. Bars represent mean ± SEM (n = 3, each replicate consists of a pool of 12 roots). Three independent experiments showing similar results were performed. (C) Fe content in shoots and roots of WT, *myb28*, *myb29* and *myb28myb29* plants grown hydroponically and subjected to a 10-days exposure to control (+Fe) or Fe-deficiency (-Fe) conditions. Bars represent mean ± SEM (n = 6-9, each replicate consists of a pool of 5 plants grown in the same hydroponic tank). Different letters indicate significant differences within each treatment (One-way ANOVA followed by Duncan’s *post hoc* test, P ≤ 0.05). Three independent experiments showing similar results were performed.

Given these observations, we next investigated whether the loss of function of MYB28 and MYB29 affects Fe accumulation. Fe content was measured in WT and mutant plants under control and Fe-deficient conditions, both in roots and shoots following 10 days of growth in control and Fe deprivation conditions (Figure 4C). Interestingly, in both conditions *myb28myb29* double mutant presented higher Fe content in roots compared to WT and single mutant plants. This later observation indicating that the observed sensitivity to Fe deficiency was not due to impaired Fe uptake but most probably to impaired Fe translocation from roots to shoots and/or distribution between plant cell types.

### myb28myb29 displays altered Fe distribution and is tolerant to Fe toxicity

To investigate Fe distribution within plant tissues, WT and *myb28myb29* plants were grown in peat under control conditions or exposed to Fe excess, to facilitate its visualization by Perls/DAB staining (Figure 5). As shown in Figure 5A, *myb28myb29* plants displayed a less pronounced damage to Fe excess compared with WT, suggesting enhanced tolerance to Fe toxicity.

**Figure 5.**
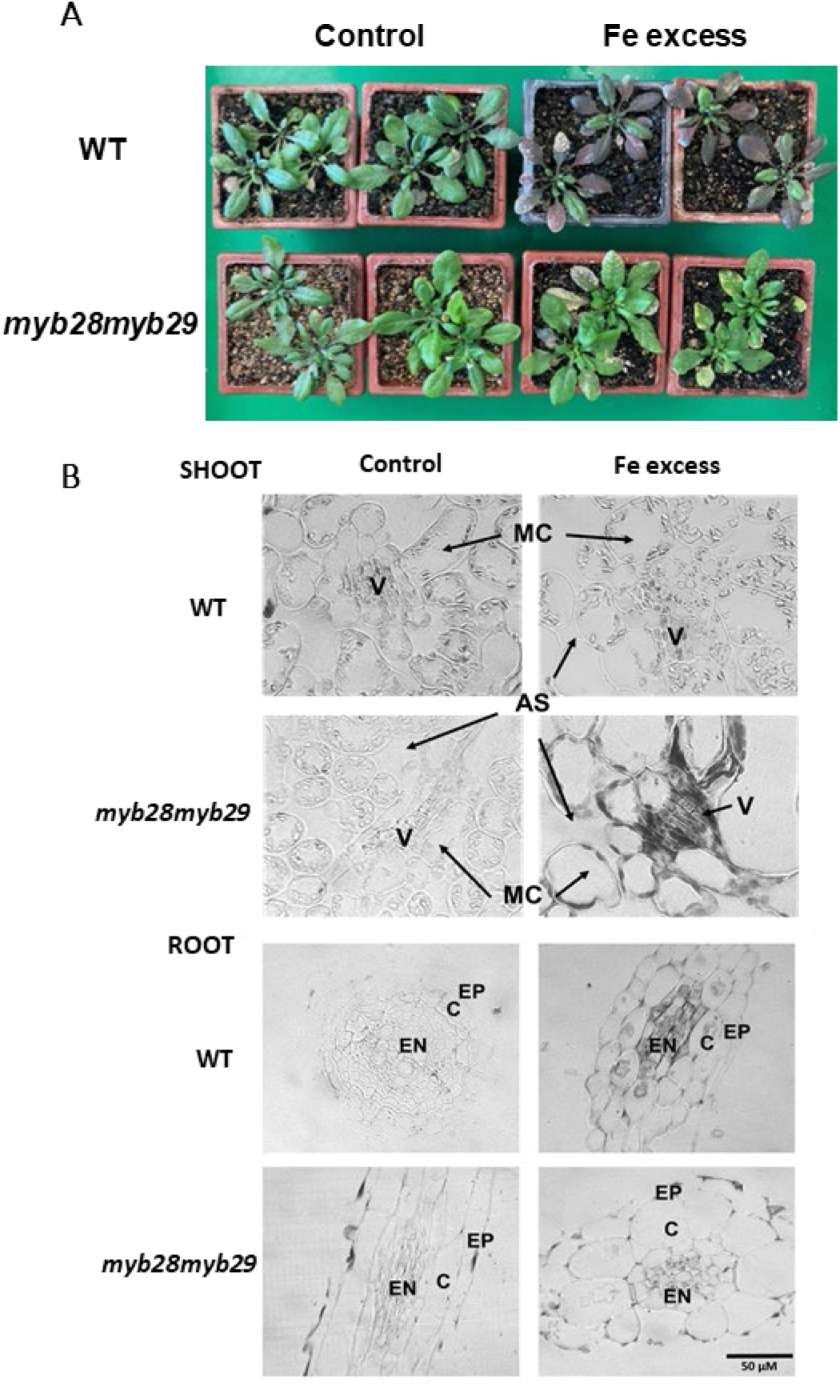
*myb28myb29* is tolerant to Fe toxicity and displays altered Fe distribution. (A) Representative images and (B) Fe distribution in shoots and roots of WT and *myb28myb29* plants grown under Fe excess. Plants were cultured in peat under control conditions (control) or after the supply of 1 g/L Fe-DTPA for two weeks (Fe excess). EP: Epidermis, C: Cortex, EN: Endodermis, V: Vasculature, MC: Mesophyll cells and AS; Apoplastic Space.

In order to localize Fe at the tissue level, leaf and root cross sections were analyzed by Perls/DAB staining (Figure 5B). In leaves exposed to Fe excess, Fe staining was more pronounced in *myb28myb29* and was particularly evident in the vasculature whereas the staining pattern in WT leaves was comparatively less intense. In roots, Fe accumulation under Fe excess was detected in the apoplast of epidermal and cortical cells of *myb28myb29*. In contrast, the vasculature showed weaker Fe staining in the mutant than in WT, in which Fe deposits were also more evident in endodermal cells (Figure 5B).

Taken together, these results suggest that in *myb28myb29* plants the alteration in Fe distribution may limit Fe toxicity and contribute to the enhanced tolerance to Fe excess, while simultaneously enhancing Fe-deficiency sensitivity.

### Fe deficiency enhances MYB28/MYB29-dependent transcriptional reprogramming

To gain further insight into the functions of MYB28 and MYB29 we performed a transcriptomic analysis (RNA-Seq) using roots from WT, *myb28*, *myb29* and *myb28myb29* plants grown under control conditions or after 10 days of Fe deficiency after transfer (Table S2). Under control conditions, the cooperative function of MYB28 and MYB29 was evident from the substantially higher number of differentially expressed genes (DEGs) detected in the double mutant compared with either single mutant (Figure 6A). A total of 372 DEGs were identified in *myb28myb29*, whereas only 47 and 53 genes were differentially expressed in *myb28* and *myb29*, respectively. Therefore, these data further support the previously reported additive roles of these TFs (Sønderby *et al*., 2010; Coleto *et al*., 2021). Under Fe deficiency, the number of DEGs greatly increased markedly in the double mutant, where 1247 genes where differentially expressed respect to WT plants, representing more than threefold increase compared with control conditions (Figure 6A). A similar trend was observed for *myb28* with 259 DEGs identified under Fe deficiency compared with 47 under control conditions. In contrast, only 18 DEGs were detected in *myb29*, consistent with the absence of a discernible Fe-deficiency phenotype in this mutant. Notably, 76% of the DEGs in *myb28* (197 of 259) were also deregulated in the double mutant. Altogether, these results reveal a cooperative function of these two TFs under Fe deficiency, while indicating that MYB28 plays the predominant role in this response.

**Figure 6.**
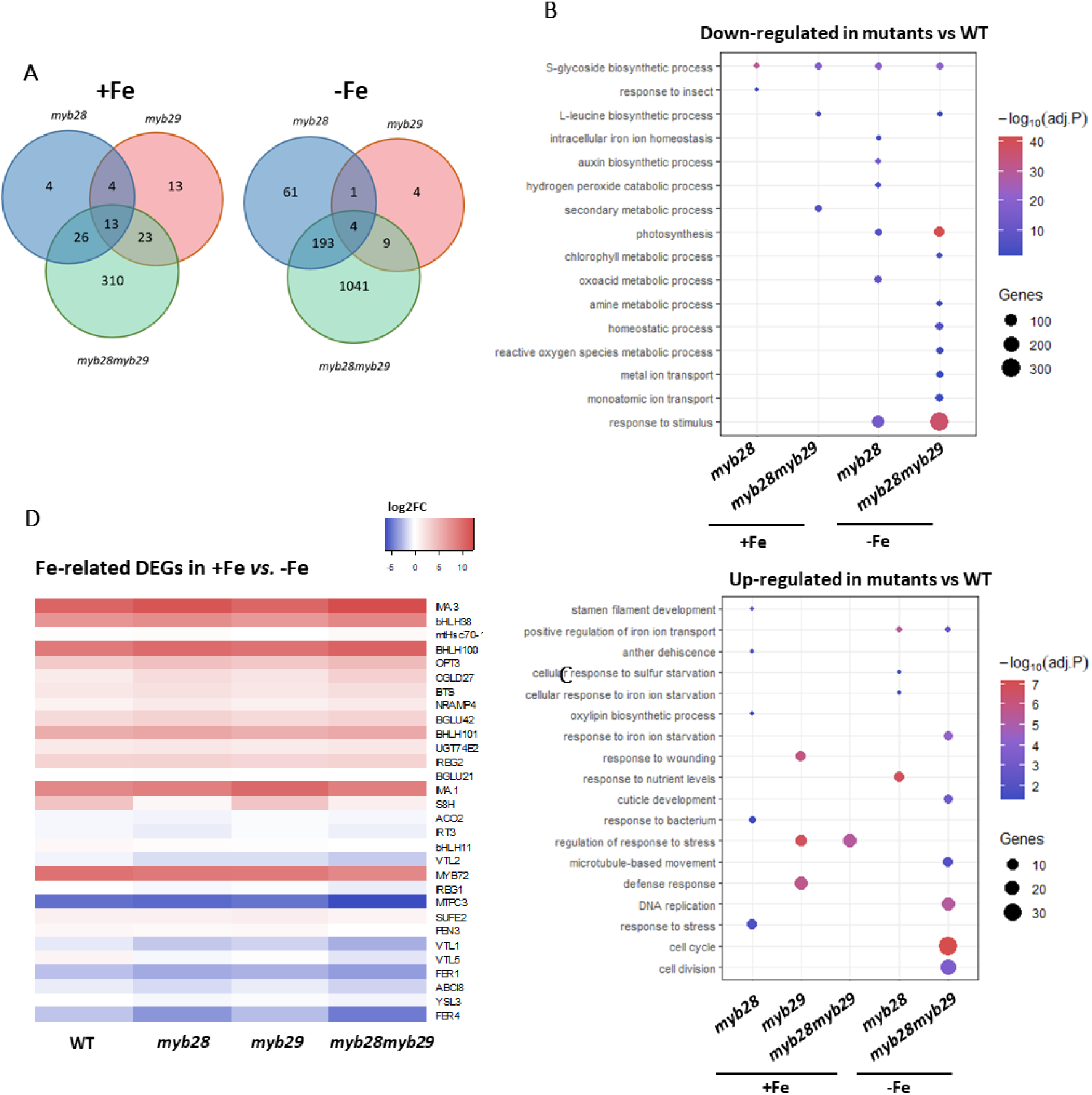
Transcriptomic analysis of WT, *myb28*, *myb29* and *myb28myb29* root in response to Fe deficiency. (A) Venn diagram and (B-C) Gene ontology (GO) enrichment or up and down-regulated genes for each genotype (P ≤ 0.05). (D) Heatmap of Fe homeostasis-related genes differentially expressed in roots under Fe deficiency conditions and in at least one WT vs mutant comparison. Plants grown *in vitro* for 7 days under control conditions and then transferred for 10 days to control (+Fe) or to Fe deficiency (-Fe). Three independent biological replicates were analysed, each one corresponding to 12 plants.

To identify biological processes affected by MYB28 and MYB29, we performed gene ontology (GO) enrichment analysis separately for up- and down-regulated genes (WT *vs* mutant). Under control conditions, enriched categories among down-regulated genes were primarily associated with glucosinolate biosynthesis and defence responses, consistent with the established functions of MYB28 and MYB29 (Figure 6B). Under Fe deficiency, additional enriched categories emerged, including terms related to redox processes and Fe homeostasis, particularly in *myb28* and, more prominently, in the *myb28myb29* double mutant. Similarly, GO analyses of up-regulated genes identified several Fe-related and stress-associated categories, further supporting a widespread deregulation of Fe-deficiency responses. Interestingly, for the double mutants GO terms related to growth, such as cell cycle and cell division were also present, in agreement with the observed biomass impairment of *myb28myb29* under Fe deficiency (Fig 6C). To examine Fe-related genes individually, we built a heatmap including those genes that were differentially expressed at least in one WT vs mutant comparison (Figure 6D). This analysis revealed a set of genes with higher induction under Fe deficiency in the mutants, consistent with the RT-qPCR results described in Figure 3, 4 and S3. The heatmap also evidenced a second set of genes exhibiting higher repression respect to WT plants, notably associated with Fe storage and/or distribution, including FERRITINS and vacuolar transporters encoding genes. Interestingly, the RNAseq analysis also underlined the expression profile of S8H, which, as expected, was greatly induced under Fe deficiency in WT and *myb29* plants but not in *myb28* and *myb28myb29*. This finding, together with RT-qPCR analysis, identifies S8H as a strong candidate downstream target of MYB28 and suggest a specific role for MYB28 in the regulation of FMC-mediated Fe homeostasis.

### MYB28 is a direct activator of S8H expression and affects FMC profile

To identify direct targets of MYB28, we initially performed chromatin immunoprecipitation (ChIP) experiments using the complemented lines expressing a MYB28-GFP fusion protein under the control of its native promoter. Although these lines successfully complemented the *myb28myb29* phenotypes (Fig 2E,F), repeated ChIP experiments yielded insufficient amounts of immunoprecipitated DNA for downstream analyses. This may reflect either low protein stability, transient or labile DNA binding, or limited chromatin occupancy. To circumvent these limitations, we performed DNA affinity purification sequencing (DAP-seq). Using this approach, we identified 398 significant MYB28-binding peaks (q=1) based on GEM parameters accounting for the characteristic bell-shaped profile of binding events, sufficient enrichment over the Halo-input signal, and the presence of motifs identified during the initial stages of GEM analysis (Table S3). On the identified peaks, 65% were located upstream of the transcription start sites (-10 Kb), while 18.1% and 14.5% were located within gene bodies and downstream of stop codons, respectively (Figure 7A). Nonetheless, 82.1% of the peaks were located within the −2 to +2 Kb region surrounding transcription start sites, indicating a strong enrichment of MYB28 in promoter-proximal regions. Motif enrichment revealed significant enrichment of an AC-element (ACC(T/A)A(C/A)(C/T) (Fig. 7B), corresponding to a previously characterized regulatory element recognized by R2R--MYB TFs (Prouse and Campbell, 2012; Kelemen et al., 2015).

**Figure 7.**
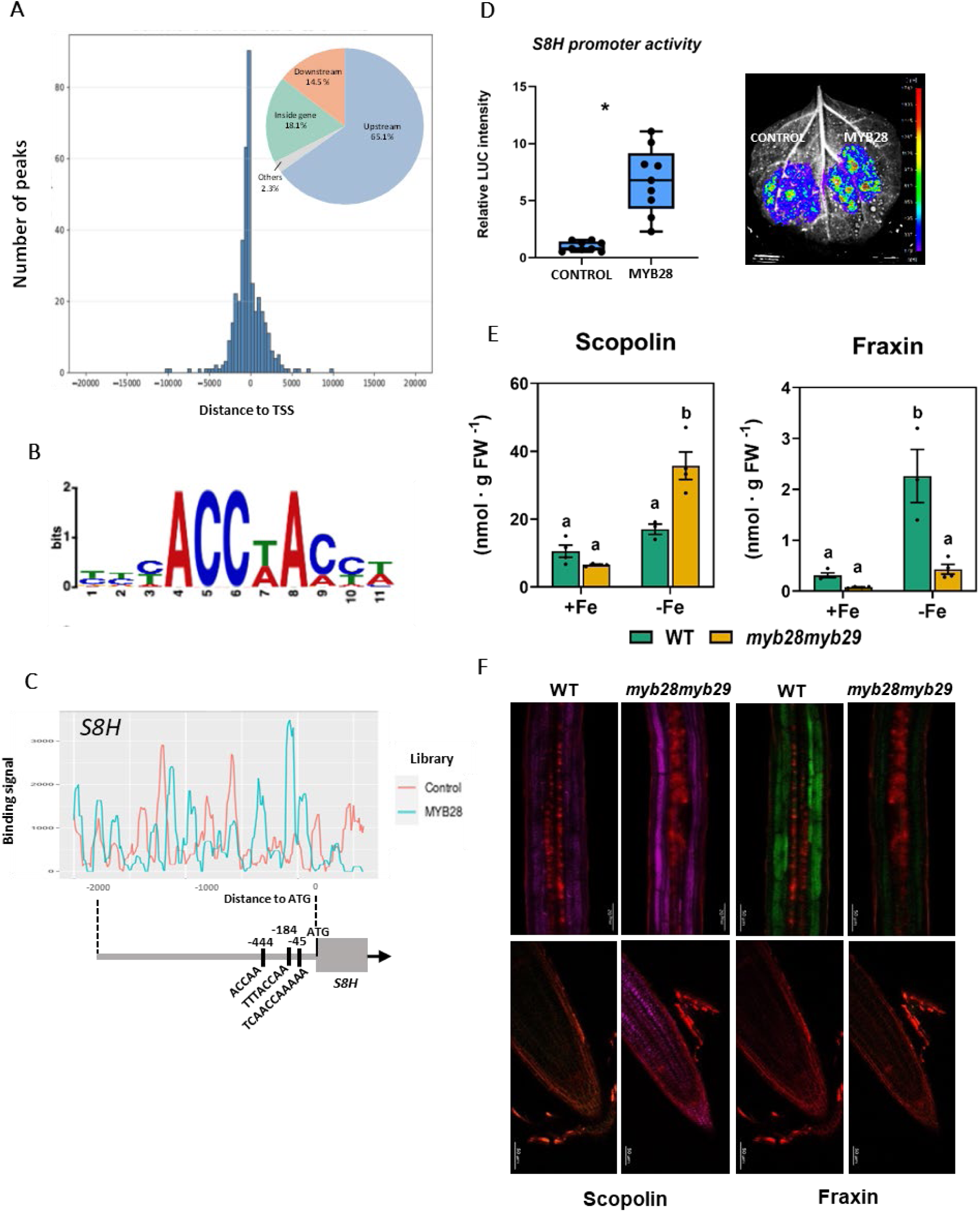
MYB28 is a direct transcriptional regulator of *S8H expression*. (A) MYB28 peak distribution regarding the distance from the transcription start site (TSS). The pie chart shows the percentage of peaks present in the upstream, downstream and inside the coding region of the nearest gene. ‘Others’ pie category accounts for peaks overlapping start or stop codon. (B) Sequence logo of MYB28 binding motif (AC-rich element). (C) MYB28 DNA-binding density plot surrounding the *S8H* transcription start site. The negative control corresponds to an input library generated using an empty GST-HALO vector. Positions of predicted MYB28 binding sites relative to the ATG start codon are indicated. (D) Activation of *S8H* promoter by MYB28 assessed via dual luciferase reporter assay in *N. benthamiana* leaves. LUCIFERASE (LUC) activity was normalized to RENILLA (REN) levels. Asterisks denote statistically significant differences (p < 0.05, t-test). (E) Scopolin and fraxin content in roots of WT and *myb28myb29* plants. Plants were grown hydroponically and subjected to a 4-day exposure to control (+Fe) or Fe-deficiency (-Fe) conditions. Bars represent means ± SEM (n=3), each replicate consists of a pool of 5 plants grown in the same hydroponic tank. Different letters indicate significant differences within each treatment (One-way ANOVA followed by Duncan’s *post hoc* test, P ≤ 0.05). (F) Visualization of scopolin and fraxin in WT and *myb28myb29* roots. Spectral images were obtained by multiphotonic microscopy (excitation wavelength: 720 nm) of seedlings grown on *in vitro* with 50 µM Fe-EDTA and then transferred 3 days under pH 7 and 100 µM FeCl_3_.

The number of significant MYB28 peaks was relatively low compared with that commonly reported for Arabidopsis TFs (O’Malley et al., 2016). This could reflect low MYB28 protein stability, weak or transient DNA-binding. Alternatively, MYB28 binding to certain genomic regions may require interacting partners to stabilize its association with DNA, consistent with the difficulties encountered in ChIP experiments. Indeed, among genes involved in aliphatic glucosinolate synthesis pathway, only *BCAT4* was associated with a significant MYB28 peak under our peak-calling criteria. However, closer inspection of the DAP-seq and Halo-input signals using a genome browser revealed additional binding profiles at several genes involved in aliphatic glucosinolate biosynthesis, as shown for instance for *CYP83A1* (Figure S4A). To validate DAP-seq results *in planta*, we employed a dual luciferase reporter assay in *Nicotiana benthamiana* and observed a strong activation of *CYP83A1* promoter in the presence of MYB28 (Figure S4B) supporting its direct transcriptional regulation by MYB28 *in planta*. Indeed, MYB28 binding to *CYP83A1*promoter had previously been reported *in vitro* by EMSA (Aarabi et al., 2016) and also by yeast on-hybrid experiments (Gigolashvili et al., 2007).

*S8H* was also not associated with a significant peak under the DAP-seq peak-calling criteria, but inspection of the binding profile revealed a clear MYB28-enriched signal compatible with a direct regulation. The putative MYB28-binding regions also contained AC-elements consistent with the binding preference identified by our motif enrichment analysis (Figure 7B). We therefore tested the ability of MYB28 to regulate *S8H* promoter using a dual luciferase reporter assay in *N. benthamiana* leaves. MYB28 strongly activated *S8H* promoter activity, further supporting *S8H* as a direct target of MYB28 (Figure 7D).

To further assess the involvement of MYB28 and MYB29 in FMC biosynthesis, we determined coumarin accumulation by HPLC and observed a significant accumulation of scopolin, the substrate of S8H, in *myb28myb29* under Fe deficiency (Figure 7E), in agreement with the lack of induction in response to Fe deficiency reported for *S8H* in this mutant. Moreover, fraxin content, the product of S8H activity was greatly reduced in *myb28myb29* mutant (Figure 7E). Finally, we took advantage of the natural fluorescence of some coumarins when exposed to UV light and further analysed their accumulation and localization in the root of seedlings grown under Fe deficiency (Figure 7F). We observed significant scopolin accumulation in cells of *myb28myb29* root cortex and endodermis, and to a lower extent in the epidermis, while the content of fraxin was greatly reduced in the same tissues. Scopolin accumulation was also observed in the apex of mutant roots (Figure 7F). Collectively, these results demonstrate that MYB28 regulates fraxin biosynthesis through activating *S8H* expression.

## Discussion

Iron deficiency is one of the most widespread nutritional stresses affecting plants, particularly in calcareous and alkaline soils where Fe solubility is severely restricted. To cope with this limitation, dicots have evolved a highly coordinated regulatory network integrating Fe sensing, transcriptional reprogramming, rhizosphere acidification, ferric reduction, metal transport and the secretion of phenolic compounds that facilitate Fe mobilization (Kobayashi *et al*., 2019; Gao and Dubos, 2021). The transcriptional network associated with the tight regulation of Fe homeostasis involves a great number of TFs, most belonging to bHLH family. However, this network expands to TFs of other families including MYB family (Gao and Dubos, 2021). For instance, MYB10 and MYB72 were shown to regulate Fe deficiency, with *myb10myb72* mutation leading to seedling mortality in plants grown in Fe-deficient alkaline soil, which could be reverted with exogenous Fe application (Palmer et al., 2013; Zamioudis et al., 2014). MYB72 was also shown to be a positive regulator of coumarin biosynthesis (Zamioudis et al., 2014). More recently, MYB8 was also shown to participate in Fe homeostasis through the activation of IRT1 transcription (Gong *et al*., 2024), whereas MYB30 interaction with BTSL1 and 2 safeguards FIT from proteosome mediated degradation, thereby resulting in enhanced FIT stability to promote the transcriptional response to Fe deficiency (Zhao *et al*., 2025).

Here we demonstrate that MYB28 and MYB29 TFs are required for an adequate adaptation to Fe deficiency and uncover an unexpected molecular connection between specialized metabolism and Fe homeostasis. Our previous work showed that the *myb28myb29* double mutant displays hypersensitivity to NH_4_^+^, a phenotype associated with alterations in Fe homeostasis (Coleto *et al*., 2021). However, it was unclear whether those alterations represented an indirect consequence of the metabolic reprogramming caused by NH_4_^+^ in *myb28myb29* plants or reflected a direct role of MYB28 and MYB29 in Fe nutrition. The present study clearly supports the latter hypothesis. For instance, loss of MYB28 consistently increased sensitivity to Fe deficiency, resulting in enhanced chlorosis, reduced root growth and impaired biomass accumulation. Although MYB29 alone produced little or no phenotype, the stronger defects observed in the double mutant, together with the transcriptomic analyses, indicate that both TFs contribute cooperatively to this response, with MYB28 assuming the predominant role. This relationship mirrors the previously established additive regulation for these two TFs in aliphatic glucosinolate synthesis (Sønderby *et al*. 2010; Coleto *et al*., 2021), suggesting that their functional redundancy extends to multiple physiological processes.

One of the most intriguing observations emerging from this work is that enhanced Fe-deficiency symptoms in *myb28* and *myb28myb29* plants cannot be explained by a lower Fe uptake. Instead, several independent observations point towards an altered perception of Fe nutritional status. The mutants displayed stronger induction of canonical Fe-deficiency marker genes, including *ILR3*, *FIT*, *bHLH38, BTS, IMA1*, *as well as FRO2* and *IRT1*. Indeed, *myb28myb29* even accumulated more Fe in roots under control or Fe deficiency conditions. These results indicate that the canonical Fe-deficiency signaling cascade remains fully operational in the absence of MYB28 and MYB29 and strongly suggest that mutant plants experience a lower physiological Fe availability despite maintaining optimal total Fe pools.

Increasing evidence indicates that total tissue Fe concentration is often a poor predictor of Fe nutritional status because only a fraction of cellular Fe is metabolically available. Several Arabidopsis mutants impaired in Fe trafficking or intracellular compartmentalization exhibit constitutive activation of Fe-deficiency responses despite accumulating Fe concentrations similar to, or higher than, WT plants (Fanara *et al*., 2022; Grant-Grant, 2022; Roschzttardtz *et al*., 2011). Such phenotypes have been interpreted as defects in Fe distribution rather than Fe acquisition. Our histochemical analyses strongly support a similar scenario. Perls/DAB staining revealed abnormal Fe accumulation around vascular tissues and in apoplastic regions of mutant leaves, whereas Fe deposits detected in root endodermal cells under Fe excess were largely absent in *myb28myb29* plants. Together with the increased tolerance of the double mutant to Fe toxicity, these observations suggest that MYB28 contributes to maintaining an appropriate tissue distribution of Fe. Consequently, mutant plants appear to perceive Fe starvation despite containing similar or higher amounts of total Fe than WT.

The transcriptomic analysis further reinforced this interpretation, since Fe deficiency dramatically amplified the transcriptional differences between WT and mutant plants, particularly in the *myb28myb29* background. Besides the expected downregulation of glucosinolate biosynthetic genes, numerous genes involved in Fe homeostasis and redox metabolism were differentially expressed. Interestingly, genes involved in Fe storage and intracellular sequestration, including *FERRITINS* and vacuolar Fe transporters, exhibited stronger repression in the mutants. Because FERRITINS constitute major intracellular Fe buffering systems that contribute to preventing oxidative damage while maintaining Fe availability (Ravet *et al*., 2009), their stronger repression may reflect an attempt to mobilize intracellular Fe reserves in response to a perceived Fe-deficient state. Likewise, altered expression of vacuolar transporters could further contribute to disturbed Fe partitioning within the cell and among tissues (Ram *et al*., 2021). Altogether, these observations support the notion that MYB28 and MYB29 influence Fe homeostasis primarily through controlling Fe utilization and distribution rather than uptake itself.

Among all differentially expressed genes identified in this study, *S8H* represents by far the most compelling candidate linking MYB28 to Fe homeostasis. *S8H* encodes SCOPOLETIN 8-HYDROXYLASE, the enzyme responsible for converting scopoletin into fraxetin, one of the catechol coumarins that plays a central role in Fe mobilization under alkaline conditions (Rajniak *et al*., 2018; Siwinska *et al*., 2018; Robe et al., 2025). Remarkably, whereas the expression of other major coumarin biosynthetic genes, including *F6’H1*, *BGLU42* and *PDR9*, remained unaffected or slightly induced in mutant plants, induction of *S8H* under Fe deficiency was virtually abolished in both *myb28* and *myb28myb29* mutants. This specificity is particularly interesting because it indicates that MYB28 does not globally regulate coumarin biosynthesis but instead controls a key metabolic step determining the production of one of the most biologically active FMCs.

The metabolic analyses provide strong support for this regulatory model. Both HPLC profiling and spectral fluorescence imaging revealed pronounced accumulation of scopolin in *myb28myb29* plants, accompanied by reduced levels of fraxin. These observations are fully consistent with the absence of induction of *SH8* expression and strongly suggest that disruption of MYB28 redirects metabolic flux towards the accumulation of S8H substrates at the expense of fraxetin biosynthesis, which contributes to explain the enhanced Fe-deficiency phenotype of *myb28* plants.

Our molecular analyses further indicate that *S8H* is likely to be a direct MYB28 target. For instance, DAP-seq analyses revealed MYB28-enriched binding signals at the S8H promoter, associated with the presence of AC-rich motifs consistent with MYB28 binding specificity. Moreover, transient dual-luciferase assays demonstrated strong activation of the *S8H* promoter by MYB28. The relatively low number of significant DAP-seq peaks compared with previous large-scale studies (ÓMalley et al., 2016) may reflect intrinsically weak or transient DNA binding by MYB28 or a requirement for additional interacting partners to stabilize its association with specific target promoters. Indeed, MYB28 has previously been shown to cooperate with members of the MYC and MYB TF families in the regulation of glucosinolate biosynthesis, raising the possibility that stable promoter occupancy *in vivo* requires additional transcriptional regulators. Regardless of the precise molecular mechanism, the convergence of transcriptomic, metabolic and promoter activation evidence data with DAP-seq binding signals and *in planta* promoter activation consistently supports *S8H* as a biologically relevant downstream target of MYB28.

The discovery that MYB28 regulates *S8H* considerably broadens the biological significance of this TF in the regulation of specialized metabolism. MYB28 has classically been viewed as a master regulator coordinating carbon allocation towards aliphatic glucosinolate biosynthesis during defense responses. Our results reveal that this regulatory function extends to the biosynthesis of metabolites required for Fe acquisition. Interestingly, MYB72 integrates Fe deficiency responses with induced systemic resistance through the regulation of coumarin secretion (Palmer *et al*., 2013; Zamioudis *et al*., 2014; Stringlis *et al*., 2018). Both MYB28 and MYB72 illustrate how metabolic reprogramming constitutes an integral component of Fe-deficiency adaptation rather than merely a secondary consequence of nutrient stress. Whether MYB28 and MYB72 function independently or converge within a broader regulatory network coordinating coumarin metabolism and the plant response to biotic stress represents an interesting question for future studies.

In summary, we propose a model in which MYB28 acts as a positive regulator of Arabidopsis response to Fe deficiency. Besides its established role in aliphatic glucosinolate biosynthesis, MYB28 directly activates *S8H* expression, promoting efficient conversion of scopoletin into fraxetin, a key FMC. Loss of MYB28 disrupts coumarin composition, alters Fe distribution and triggers exaggerated Fe-deficiency signaling despite largely unaltered total Fe accumulation. Our work therefore identifies MYB28 as a previously unrecognized component of the Fe homeostasis network and uncovers a novel signaling link between specialized metabolism and mineral nutrition. These findings considerably expand the biological functions of MYB28 and provide new insight into how transcriptional regulation coordinates metabolic pathways required for plant adaptation to nutrient-limiting environments.

## Methods

### Plant material and growth conditions

The plant material used in this study was wild-type (WT) *Arabidopsis thaliana* L. Col-0 ecotype and mutant lines derived from the Col-0 background. The mutant lines were *myb28*, *myb29*, and *myb28myb29* (Li et al., 2013) and *s8h-2* (Siwinska *et al*., 2018). Seeds were sterilized with 70% (v/v) ethanol for 30 sec and 35% (v/v) sodium hypochlorite for 7 min, followed by a rinsing with sterile deionized water.

For hydroponic culture, we followed the protocol of Choi et al. (2025). Briefly, seeds were placed in PCR tubes, used as support for seed germination, filled with 0.8 % (w/v) agar solidified Hoagland medium [5 mM Ca(NO₃)₂, 5 mM KNO₃, 1 mM KH₂PO₄, 1 mM MgSO₄, 50 μM MnSO₄, 50 μM H₃BO₃, 25 μM Fe³⁺-EDTA, 15 μM ZnSO₄, 3 μM Na₂MoO₄, 2.5 μM KI, 0.05 μM CoCl₂, 0.05 μM CuSO₄ (pH 5.7)]. After 7 days, seedlings were transferred to 0.5 L plastic tanks filled with liquid Hoagland solution at pH 5.7 (2.5 mM MES) or pH 7.5 (2.5 mM MOPS). Five plants were grown per tank, and nutrient solution was renewed weekly. After 4 weeks, tanks were divided in two groups, control conditions and Fe deficiency, where Fe was no longer added to the solution. Experiments were carried out in a growth chamber with 8/16 h light (150 μmol photons m^-2^ s^-1^)/dark photoperiod, 22/18 °C and 60/70 % relative humidity.

For *in vitro* plant culture, surface-sterilized seeds were placed in 12 x 12 cm square Petri dishes, containing a modified Murashige & Skoog (MS) growth medium. The medium composition was 5 mM KCl, 2.5 mM Ca(NO_3_)_2_, 2.25 mM CaCl_2_, 1.25 mM KH_2_PO_4_, 0.75 mM MgSO_4_, 100 µM MnSO_4_, 100 µM H_3_BO_3_, 85 µM Na_2_EDTA, 30 µM ZnSO_4_, 5 µM KI, 0.1 µM CuSO_4_, 0.1 µM Na_2_MoO_4_, 0.5 % sucrose and 1.5% (w/v) agar (A5431, Sigma-Aldrich). The medium was buffered with 2.5 mM MES, and the pH adjusted to 5.7. Under control conditions, Fe was supplied as 100 μM FeSO₄. Plates were kept at 4 °C and in darkness for 2 days and then placed vertically in a growth chamber under the following conditions: 22 °C and 60 % humidity during a 14 h light period (150 μmol photons · m^-2^ · s^-1^), and 18°C and 70% humidity during a 10 h of dark period. At the end of the growth period, plant images were acquired using a scanner (EPSON expression 10.000 XL). Root length was measured using FIJI software v2.14.0.

For experiments in peat, plants were grown in 7 x 7 x 8 cm pots. For Fe limitation experiments, 0.5% (w/w) CaO was incorporated to induce low Fe available conditions (Long et al., 2010). Five seeds were sown per pot. The pots were maintained in a growth chamber during 4 weeks under controlled conditions: 22°C and 60% humidity with 14 h of light (150 μmol photons · m^-2^ · s^-1^), followed by 18°C and 70% humidity for 10 h of darkness. For Fe toxicity experiments, after one week in peat, plants were watered once a week with a solution of 0.1 % (v/v) Fe-DTPA (Sprint 330, BASF) until toxicity symptoms became evident.

For all experiments plants were harvested 2 h after the onset of the photoperiod. Eventually, biomass of shoots and/or roots was measured and the plant material immediately frozen in liquid nitrogen and stored at −80 °C until use.

### Generation of MYB28 complemented lines

To generate the *myb28myb29 pMYB28:gMYB28:GFP* complemented lines, a region starting 2 kb upstream of the start codon and the whole *MYB28* gene locus was amplified from Col-0 genomic DNA using the Phusion plus Taq polymerase (Thermo Scientific) and specific primers (Table S1). The resulting 3,324 bp fragment was cloned into *pDONR207* and recombined into the binary vector *pGWB4*, using the Gateway Cloning System (Invitrogen). Subsequently, the construct was introduced into Arabidopsis *myb28myb29* plants by *Agrobacterium tumefaciens*–mediated floral dip transformation.

### Chlorophyll quantification

The content of chlorophyll was determined as described by Chazaux et al. (2022). Briefly, total chlorophyll was extracted with 80% (v/v) acetone solution (10 µl per milligram of leaf fresh weight) using a tissue homogenizer (MM400, Retsch GmbH). The homogenates were centrifuged at 16,000 *g* for 20 min at 4 °C. After centrifugation, the supernatant was recovered and diluted 10 times with 80% acetone. To determine chlorophyll, 300 μL of the extract were placed in a 96-well microplate and the absorbance was measured at 645 and 663 nm.

### Total iron quantification

The total content of mineral Fe was quantified from 25-50 mg of lyophilized root and shoot powder. Plant tissue was digested in a microwave (MARS6, CEM) in 10 mL of HNO_3_ for 15 min at 180 °C. After filtration through MN 640 W paper, Fe content was determined by inductively coupled plasma mass spectrometry (ICP-AES Optima 8300).

### Perls staining and DAB intensification

Fe staining was conducted following the methodology described in Roschzttardtz et al. (2009). Briefly, leaves and roots of the Fe toxicity experiment were fixed in a solution of 2% (w/v) paraformaldehyde in a 100 mM sodium phosphate buffer (pH 7) for 1 h under vacuum and incubated overnight at room temperature. Samples were dehydrated with a serial bath of ethanol, 50, 60, 70, 80, 90, 95, and 100% (v/v) for 1 h per bath. Dehydrated embryos were incubated with ethanol/butanol overnight and then incubated again in butanol 100% overnight. A final incubation with butanol/resin overnight at room temperature was performed. Afterwards, samples were embedded in Technovit 7100 resin (Kulzer) and 3 μm thin sections were prepared.

These sections were placed on glass slides and incubated under vacuum for 45 min in Perls stain solution (2% HCl (v/v), 2% potassium ferrocyanide (v/v)). After washing with distilled water, sections were incubated in a methanol solution containing 10 mM NaN_3_ and 0.3% (v/v) H_2_O_2_ for 1 h and washed with 100 mM Na-phosphate buffer, pH 7.4. For the intensification reaction, sections were further incubated for 10 min in 100 mM Na-phosphate buffer, pH 7.4, solution containing 0.025% (w/v) DAB hydrate, 0.005% (v/v) H_2_O_2_, and 0.005% (w/v) CoCl_2_·2H_2_O. The reaction was stopped by rinsing with distilled water. Sections from at least 3 independent plants were observed with a microscope Eclipse 80i (Nikon) and images acquired with the camera Nikon Digital Sight DS-5M.

### Gene expression analysis

Total RNA was extracted from 15 mg of frozen root tissue using the Mini Total RNA Kit for Plants (IBI Scientific), which includes DNase treatment. For expression analysis by quantitative real-time PCR (qPCR), 1 μg of RNA was retrotranscribed to cDNA using the PrimeScriptTM RT kit (Takara-Bio Inc., Otsu, Shiga, Japan). PCR was performed in a Step One Plus Real Time PCR System (Applied Biosystems). Each reaction consisted on 7.5 µl of SYBR® Premix Ex Taq™ II kit (Takara Bio Inc.), 0.3 µl of ROX, 0.5 µl (10 µM) of each gene-specific primer (Table S1) and 2 µL of cDNA diluted 1:10 in a 15 μL reaction volume. We employed a two-step PCR program that consisted of an initial denaturation and activation of Taq polymerase (95 °C for 4 min), followed by 40 cycles of amplification and quantification (94 °C for 15 s, 60 °C for 1 min). The protocol was finished with a melting curve (60–95 °C with one fluorescence reading every 0.3 °C). Relative gene expression was calculated as the ΔCp between each gene and the average of the housekeeping genes *SAND* (At2g28390) and *ϐ-TUBULIN* (At5g44340).

For RNA-seq analysis, mRNA was purified using poly(T)oligo-attached magnetic beads. Libraries were generated with the NEBNext® Ultra™ RNA Library Prep Kit for Illumina® (NEB, USA) and sequenced, generating paired-end reads on an Illumina platform. Library preparation and sequencing were outsourced to Novogene (UK). The sequencing reads were aligned to Arabidopsis reference genome TAIR 10, using HISAT2 (Mortazavi *et al*., 2008). Transcripts were quantified as fragments per kilobase of transcript per million mapped reads (FPKM) with HTseq (Anders *et al*., 2015). Differential expression was assessed using DESeq2 (Love et al., 2014) and p-values were adjusted using the Benjamini and Hochberg’s approach for controlling the false discovery rate (FDR). Genes with an adjusted P-value <0.05 and fold change>1.5 were assigned as differentially expressed. Gene ontology (GO) enrichment analysis was conducted using g:Profiler2 (Kolberg et al., 2023) with default parameters. To reduce redundancy, we focused on the representative (highlighted) GO terms identified by g:Profiler. Three independent biological replicates per genotype and treatment were analysed (each replicate corresponding to a pool of 12 individual plants).

### DNA affinity purification sequencing (DAP-seq)

Plasmid preparation: *MYB28* CDS (1.1 kb) was amplified by PCR using the Phusion plus Taq polymerase (Thermo Scientific), cloned into *pDONR207* and recombined into the Gateway-compatible *pIX-HALO* expression vector (Bartlett et al., 2017), which contains an N-terminal HaloTag affinity sequence. Primers are given in Table S1. Arabidopsis genomic DNA extraction was performed as indicated in Thomas et al. (1993). DAP-seq was conducted as in Bartlett et al. (2017), with modifications as described below:

Genomic DNA library preparation: DNA was sonicated to obtain 200 bp fragments on a Covaris Focus-ultrasonicator instrument. Fragments were selected by AMPure XP beads for removing small fragments (<100 bp). Beads with correct size fragments were washed twice with 80% ethanol, then resuspended in elution buffer after ethanol evaporation. Selected fragments were processed in the following three steps, including (i) end repair, (ii) adding A-tail and (iii) adapter ligation. Two cleaning events, following the instruction of the QIAquick PCR Purification Kit (Cat No./ID: 28104), were required among these three steps. After the adapter ligation, fragments fused with adapters, were purified and washed with AMPure XP beads and 80% ethanol for removing unattached adapters. Resuspended ligated fragments were quantified by Qubit and were confirmed by Bioanalyzer analysis.

Protein preparation: The MYB28-pHALO recombinant plasmid and an empty pIXHALO plasmid (which produces only the HALO peptide when translated) were *in vitro* translated with TNT Coupled Reticulocyte Lysate System (Promega). Three replicates were performed. The pIX-HALO plasmid was used as the ‘input’ control.

DAP-seq library preparation: Washed Halo-Tag beads were mixed with the *in vitro* translated proteins to pull down the HALO-MYB28 protein. Each pulled-down protein reaction was mixed with 500 ng genomic DNA libraries for MYB28 binding. The bound DNA was washed by PBST for six times and then released from Halo-Tag beads by denaturation at high temperature (98°C). The eluted DNA was enriched by PCR amplification to generate the sequencing libraries. The enriched DAP-seq libraries were purified and washed with AMPure XP beads and 80% ethanol for removing indexes. Resuspended DAP-seq libraries were quantified by Qubit and were confirmed by Bioanalyzer analysis.

DAP-seq library sequencing: DAP-seq libraries were sequenced on an Illumina NextSeq 550 (sequencing of libraries was set at 40 million and 1×75bp single-end reads). As a negative control, the ‘input’ (*pIX-HALO* expression vector without any insertion of opening reading frame) was accounted for possible non-specific DNA binding in the reaction mixture, and copy number variations at specific genomic loci. DAP-seq reads were mapped to the *Arabidopsis thaliana* TAIR 10 reference genome using bowtie2 (Langmead and Salzberg, 2012), version 2.0-beta7, with default parameters and post-processing to remove reads that have MAPQ scores lower than 30. Peak detection was performed using GEM peak caller (Guo et al., 2012) version 3.4 with the TAIR10 genome assembly using the following parameters: ‘-q 2 -t 1 -k_min 6 - kmax 20 -k_seqs 600 -k_neg_dinu_shuffle’. The replicates were analysed as multi-replicates with the GEM replicate mode. Peak summits called by GEM were associated with the closest gene model in the Araport11 annotation file using the BioConductor package ChIPpeakAnno (Zhu et al., 2010) with default parameters (i.e. NearestLocation). De novo motif discovery was performed by retrieving 200-bp sequences centred at GEM-identified binding events for the significant peaks and running STREME in MEME suit 5.5.3 with most default parameters. Motifs were screened using the Tomtom tool in MEME to compare to previously published motifs found in the JASPAR database (subdomain plants and Arabidopsis).

### Coumarins visualization and quantification

Coumarins were analysed as described in Robe et al., (2021b) and Watanabe et al., (2025). Quantification was performed by high-performance liquid chromatography (HPLC) using 70 mg of frozen root powder was extracted in 700 μL 80% methanol (v/v) and filtered through a 0.45 μm PVDF filter. Coumarin visualization was acquired from 7-d-old seedlings grown on 0.5X MS medium with 50 µM Fe-EDTA for 4 days and then, transferred 3 additional days to half strength MS with 100 µM FeCl_3_ pH 7 or 50 µM Fe-EDTA pH 5.7. Root cell walls were stained with a solution of 10 µg ml^−1^ propidium iodide (PI) for 10 min and fraxin and scopolin were imaged using their autofluorescence properties with an LSM 880 multiphotonmicroscope (Zeiss, Oberkochen, Germany).

### Dual luciferase assay

Promoter regions of *S8H* (2 kb) and *CYP83A1* (1.72 kb) respectively, were cloned into the pPGWL7.0 reporter vector (Karimi et al., 2002), using the Gateway Cloning System (Invitrogen) to control the expression of the firefly luciferase gene (*LUC*). Similarly, the *MYB28* CDS was cloned into *pBin19-35S-GW-HA* (Marino et al., 2013). The *S8Hpro:LUC*, *CYP83A1pro:LUC* reporter vectors, *35S:MYB28* effector vector, and the *Renilla reniformis* (*REN*) reference vector (*pK7WG2 – 35S: REN*) were transferred to *A. tumefaciens* C58C1. Agrobacterium harbouring the vector *pCH32-35S:p19*, which expresses the silencing suppressor p19 of tomato bushy stunt virus, was also used. Overnight *A. tumefaciens* cultures expressing the constructs of interest were harvested by centrifugation (3000 *g,* 15 min). Cells were resuspended in induction buffer (10 mM MgCl_2_, 10 mM MES, pH 5.6, and 150 mM acetosyringone) to an OD600 of 0.2. After 3 h at 22 °C, cells were mixed in 1:1 ratio and infiltrated into leaves of 4-week-old *Nicotiana benthamiana* plants. Three days after infiltration, one cm diameter leaf discs were harvested, immediately placed in liquid nitrogen and stored at –80 °C. Dual Luciferase Assay was carried out as using the Dual-Luciferase® Reporter (DLR™) Assay System following the manufacturer instructions (Promega). Results were obtained from the analysis of 1 leaf disk per agroinfiltrated plant (9 plants per condition). LUC and REN luminescence was acquired and detected using a Biotek SYNERGY HTX multi-mode reader and the data were presented as the ratio of the two measurements relativized to control condition. For images, leaves were sprayed with 1 mM D-Luciferin potassium salt, incubated in the dark for 2 minutes and imaged using a cooled CCD camera (NightOwl II LB 983 NC-100; Berthold Technologies, Bad Wildbad, Germany).

### Statistical analysis

Data were analysed using SPSS software (version 26.0, IBM Corp., Chicago, IL, USA). Normality was assessed using the Kolmogorov–Smirnov test, while homogeneity of variances was evaluated using Levene’s test. Pairwise comparisons were performed using Student’s t-test. For multiple comparisons, a one-way ANOVA followed by Duncańs *post-hoc* test was used to compare means between treatments.

## Data availability

RNA-seq and DAP-seq data have been deposited in GEO under the following IDs, respectively: GSE341148 and GSE346451.

Accessions numbers for genes can be found in Table S1.

## Author contributions

DM conceived the project. DM and CD funding acquisition. AJMP, IC, JAUG, AR, HR, JM, CD and DM designed experiments. AJMP, IC, JAUG, AR, DT, SW, DSG, MBT, HR, JM, performed experiments. AJMP, IC, JAUG, AR, DT, SW, AS, HR, JM, JTM, CD and DM analysed data. DM and CD wrote the initial manuscript, with inputs from other authors.

## Acknowledgements

The authors thank for technical and human support provided by SGIker (EHU/ ERDF, EU). We thank Dr Carine Alcon and Matthieu Déjean for technical assistance and expertise for microscope observations and the imaging facility MRI, member of the national infrastructure France-BioImaging supported by the French National Research Agency (ANR-10-INBS-04, ‘Investments for the future’). We thank Prof. Masami Hirai (RIKEN) for kindly providing seeds for *myb28*, *myb29* and *myb28myb29* mutants.

## Declaration of Generative AI and AI-assisted technologies in the writing process

During the preparation of this work the authors used ChatGPT to improve the grammar and readability of the manuscript. After using this tool/service, the authors reviewed and edited the content as needed and take full responsibility for the content of the publication.

---

**Supplementary Figure S1.**
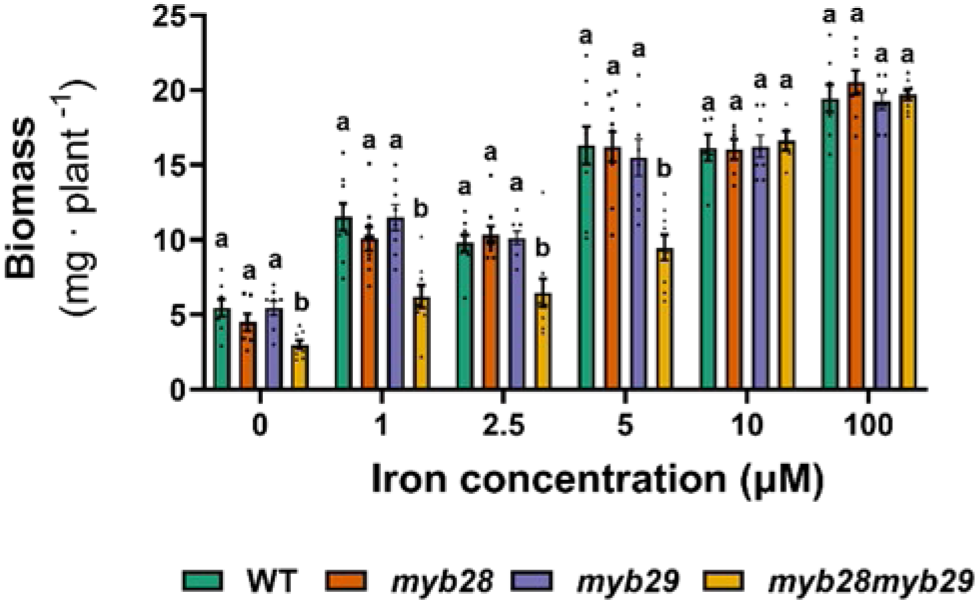
Biomass of WT, *myb28*, *myb29* and *myb28myb29* plants grown *in vitro*. Bars represent mean ± SEM (n = 7-10, each replicate consists of a pool of 10 plants). Different letters indicate significant differences within each treatment (One-way ANOVA followed by Duncan’s *post hoc* test, P ≤ 0.05). Three independent experiments showing similar results were performed.

**Supplementary Figure S2.**
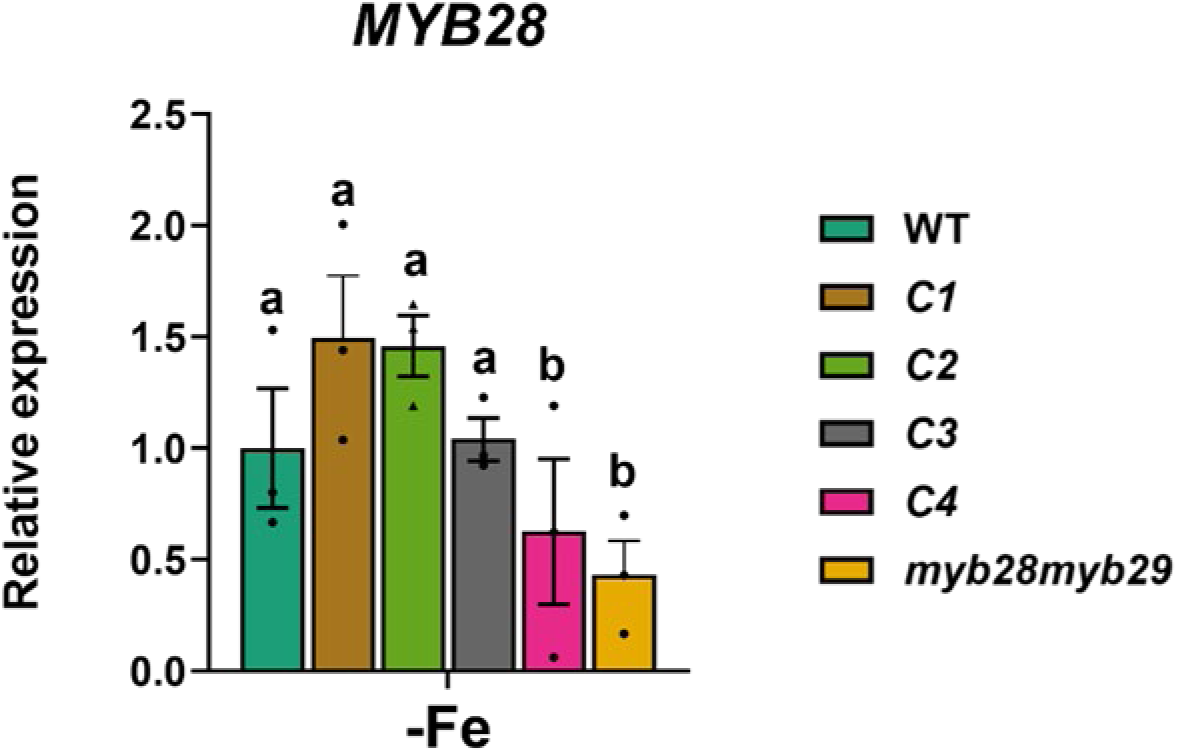
*MYB28* expression in roots of *myb28myb29* plants complemented with MYB28. Plants (*pMYB28:gMYB28:GFP* #1, C1; *pMYB28:gMYB28:GFP* #2, C2; *pMYB28:gMYB28:GFP* #3, C3 and *pMYB28:gMYB28:GFP* #4, C4) were cultured *in vitro* for 10 days under Fe deficiency conditions (-Fe). Bars represent mean ± SEM (n = 3, each replicate consists of a pool of 10 roots). Different letters indicate significant differences within each treatment (One-way ANOVA followed by Duncan’s *post hoc* test, P ≤ 0.05). Three independent experiments showing similar results were performed.

**Supplementary Figure S3.**
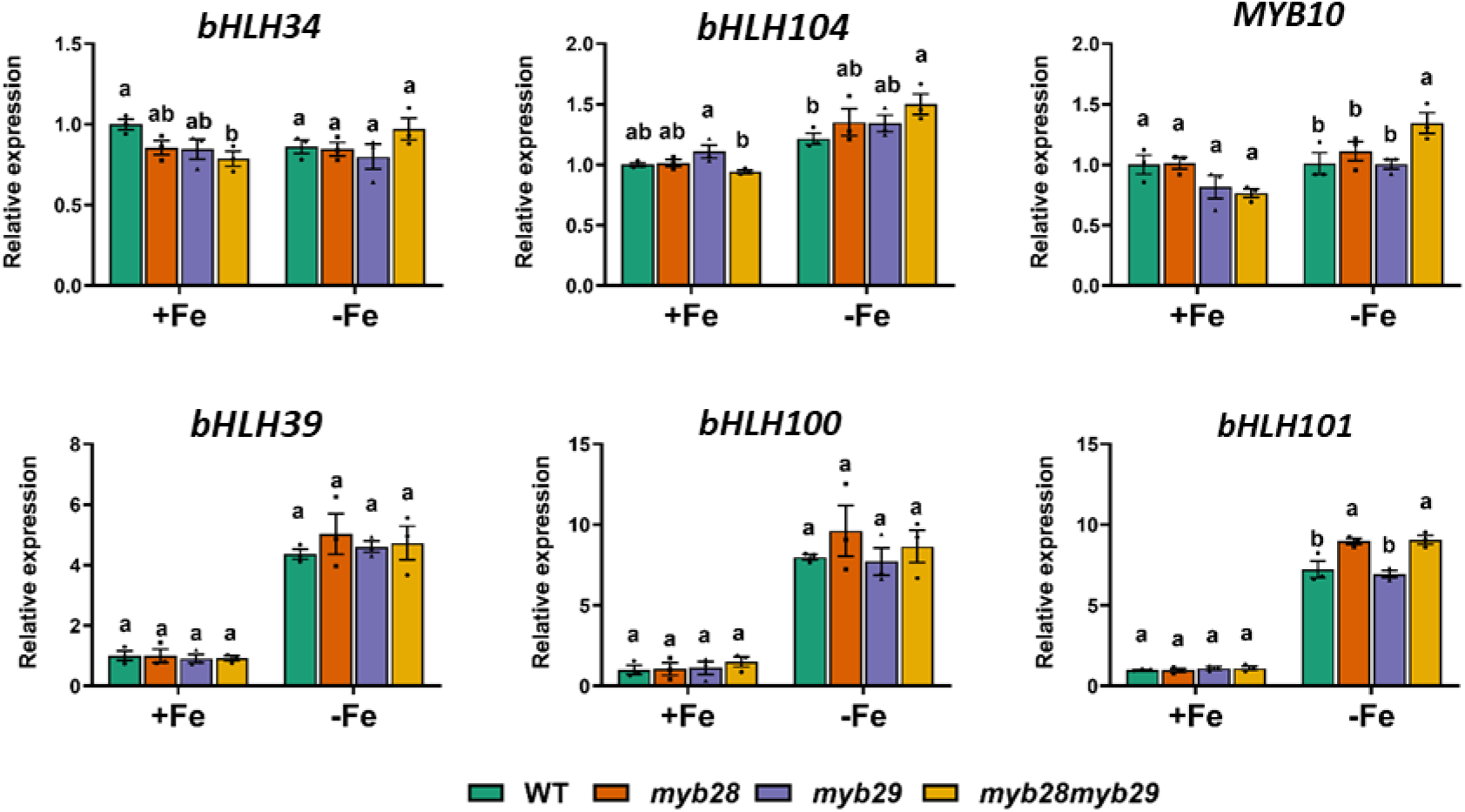
Expression of Fe homeostasis-related genes. WT, *myb28*, *myb29* and *myb28myb29* plants were grown *in vitro* under control (+Fe) or Fe deficiency (-Fe) conditions for 15 days. Bars represent mean ± SEM (n = 3, each replicate consists of a pool of 12 roots). Different letters indicate significant differences within each treatment (One-way ANOVA followed by Duncan’s *post hoc* test, P ≤ 0.05). Three independent experiments showing similar results were performed.

**Supplementary Figure S4.**
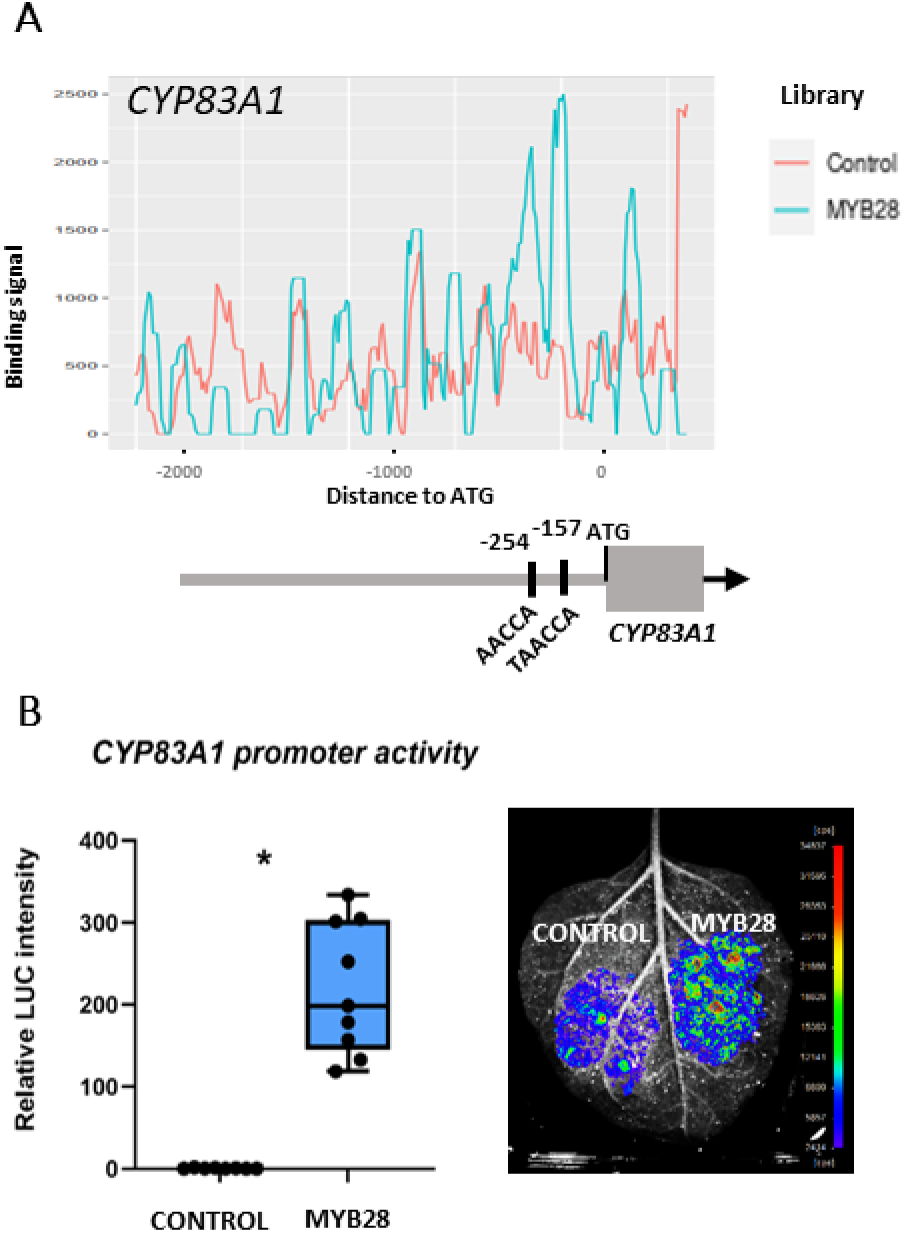
MYB28 activates CYP83A1 promoter activity. (A) MYB28 DNA-binding density plot surrounding the *CYP83A1* transcription start site. The negative control corresponds to an input library generated using an empty GST-HALO vector. Positions of predicted MYB28 binding sites relative to the ATG start codon are indicated. (B) Activation of *CYP83A1* promoter by MYB28 assessed via dual luciferase reporter assay in *N. benthamiana* leaves. LUCIFERASE (LUC) activity was normalized to RENILLA (REN) levels. Asterisks denote statistically significant differences (p < 0.05, t-test).

**Supplementary Table S1.**
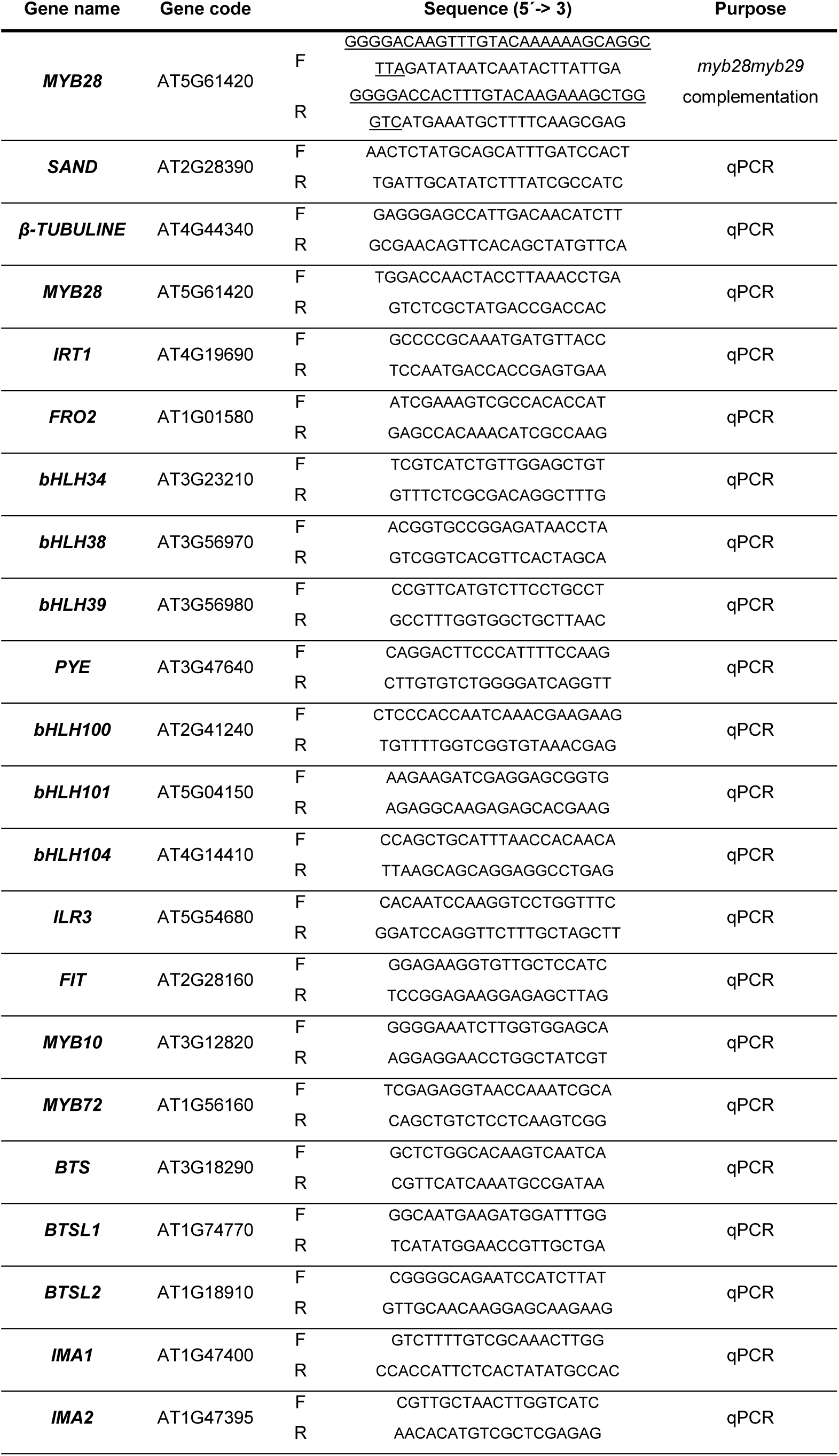
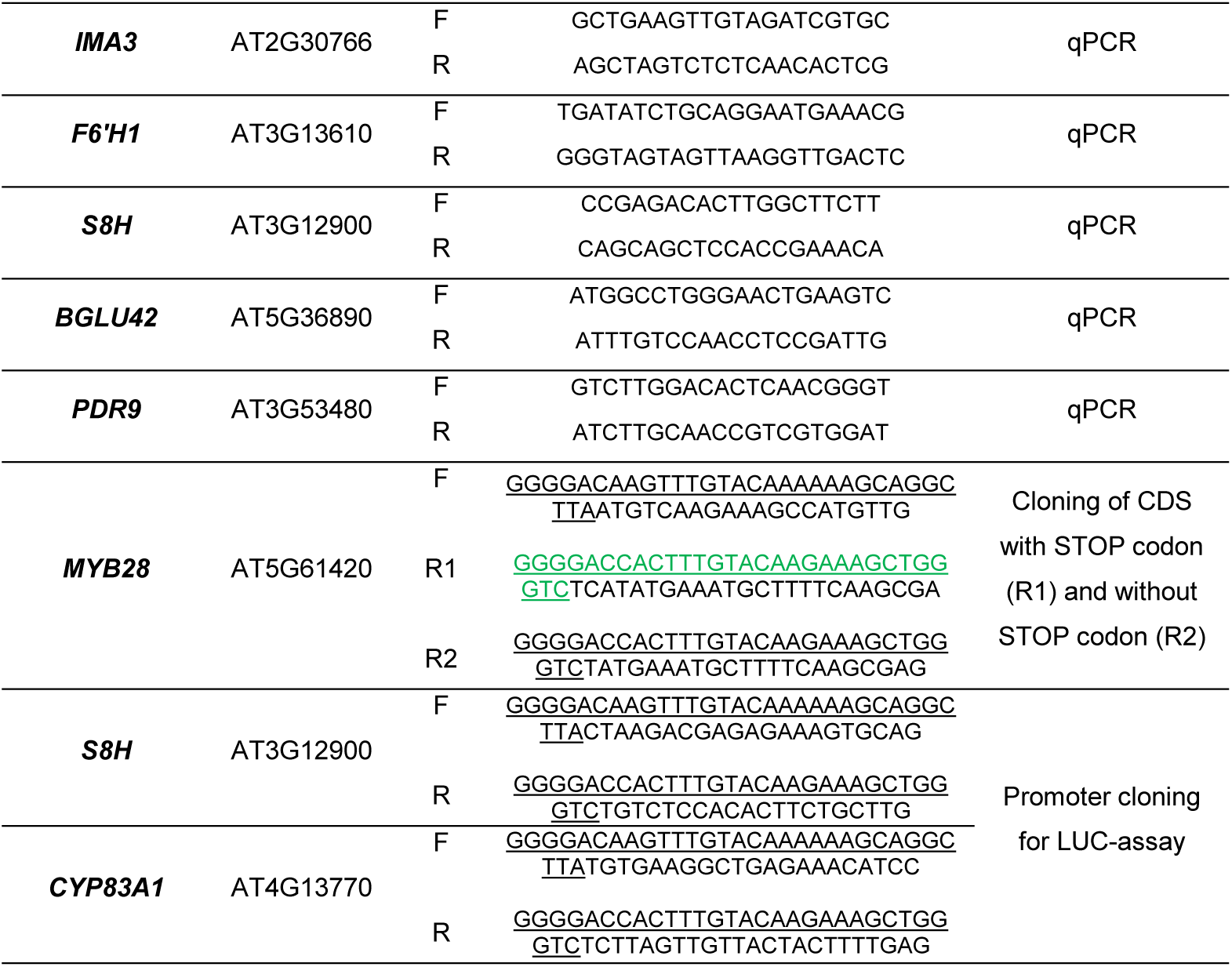
List of primers used. Recombination flanking regions for Gateway cloning technology are underlined.

**Supplementary Table S2.** Differentially expressed genes in response to Fe deficiency in WT, *myb28*, *myb29* and *myb28myb29* plants.

**Supplementary Table S3.** List of MYB28-binding peaks detected by DAP-seq analysis.

